# *Drosophila* AHR limits tumor growth and stem cell proliferation in the intestine

**DOI:** 10.1101/2023.05.17.538815

**Authors:** Minghua Tsai, Jiawei Sun, Cyrille Alexandre, Michael Shapiro, Adrien Franchet, Ying Li, Alex P. Gould, Jean-Paul Vincent, Brigitta Stockinger, Nicola Laura Diny

**Affiliations:** The Francis Crick Institute, NW1 1AT London, UK

## Abstract

The aryl hydrocarbon receptor (AHR) plays important roles in intestinal homeostasis, limiting tumour growth and promoting differentiation in the intestinal epithelium. Spineless, the *Drosophila* homolog of AHR, has only been studied in the context of development but not in the adult intestine. Here, we show that *spineless* is upregulated in the adult intestinal epithelium after infection with *Pseudomonas entomophila* (*P*.*e*.). Spineless knockdown increased stem cell proliferation following infection-induced injury. Spineless overexpression limited intestinal stem cell proliferation and reduced survival after infection. In two tumour models, using either *Notch* RNAi or constitutively active Yorkie, Spineless suppressed tumour growth and doubled the lifespan of tumour-bearing flies. At the transcriptional level it reversed the gene expression changes induced in Yorkie tumours, counteracting cell proliferation and altered metabolism. These findings demonstrate a new role for Spineless in the adult *Drosophila* midgut and highlight the evolutionarily conserved functions of AHR/Spineless in the control of proliferation and differentiation of the intestinal epithelium.

## Introduction

The aryl hydrocarbon receptor (AHR) is a ligand-activated transcription factor with barrier-protective roles in the intestine^1^. AHR is an environmental sensor of the basic helix-loop-helix Per-Arnt-Sims (bHLH-PAS) family that binds ligands derived from the diet, microbial metabolism or industrial sources. Ligand binding leads to release of AHR from its chaperone complex and nuclear translocation^2^. AHR then dimerizes with AHR nuclear translocator (ARNT) for DNA binding at canonical binding sites. AHR is widely expressed in many intestinal immune cells, stromal cells and the intestinal epithelium^3^, where it is important in the anti-bacterial defence and in limiting tumour growth^1^. Ablation of AHR in the intestinal epithelium of mice leads to increased susceptibility to infection with *Citrobacter rodentium* and increased malignant transformation in an AOM-DSS model^4^. AHR is needed to end the regenerative response of the intestinal epithelium after injury to allow the epithelial barrier to return to its mature state^5^. Treatment of mice with AHR ligand-rich diet was shown to be beneficial in tumour models and epithelial healing after injury with DSS^4,5^.

Spineless is the closest *Drosophila* homolog to AHR and binds the same DNA sequence^6,7^. Amino-acid identity between AHR and Spineless is 41% overall but substantially higher in the PAS domains and reaches 70% in the DNA-binding site^8^. The lowest similarity is found in the ligand-binding domain. This is in line with the idea that Spineless is a ligand-independent transcription factor that cannot bind prototypic AHR ligands like dioxin and does not require ligands for nuclear translocation^9,10^. Moreover, *Drosophila* is unaffected by dioxin^11^. Akin to the AHR-ARNT dimer in vertebrates, Spineless forms a heterodimer with the bHLH-PAS family member Tango and this heterodimer then translocates to the nucleus to bind to dioxin-response elements^7^. Another line of evidence for strong evolutionary conservation between these homologs comes from a study showing that AHR could rescue the developmental phenotypes of Spineless mutants in *Drosophila*^11^. The authors also demonstrated that AHR/Spineless functions in *Drosophila* are highly dependent on gene dosage of Tango or ARNT and that dioxin treatment could enhance AHR functions in murine AHR-transgenic *Drosophila*. Thus, *Drosophila* Spineless might be a useful model to study evolutionarily conserved AHR functions.

The functions of Spineless have been studied extensively in *Drosophila* development. Spineless was first identified for controlling antenna development, with mutants causing the aristapedia phenotype^12^. Spineless has since been shown to function together with Tango to control antennal identity and the development of tarsal segments of the leg^7,8,13,14^. Spineless also plays important roles in regulating dendrite morphology in neurons^15^, the development of sternopleural bristles^16^, and photoreceptor specification in the retina^17-21^. Few studies have focused on the function of Spineless in adult flies and the role of Spineless in the adult intestine has not been studied.

The *Drosophila* midgut consists of a single layer of epithelial cells surrounded by a basement membrane and visceral muscle. Similar to the mammalian intestine, the epithelium regenerates from intestinal stem cells (ISC), which give rise to transient enteroblasts (EB) and further differentiate into mature absorptive enterocytes^22^. The *Drosophila* intestine also contains a secretory lineage, the enteroendocrine cells. ISC are characterized by expression of Escargot (Esg) and Delta (Dl) and suppression of Notch (N) signalling for their maintenance^23,24^. Upon asymmetric division and differentiation into EB, cells downregulate Delta, activate Notch signalling and induce Suppressor of Hairless (Su(H)). Differentiation into Pou domain protein 1 (Pdm1)-positive enterocytes is driven by Jak/Stat signalling, Notch activation and a downregulation of Esg. Prospero (Pros) is induced during differentiation into enteroendocrine cells. A range of local, systemic, and environmental stimuli are integrated through multiple signalling pathways (including Notch, Jak/Stat, Egfr and Hippo) to govern ISC proliferation and differentiation and to maintain the epithelial barrier and its function^22^. Many of these pathways have also been shown to interact with AHR to regulate stem cell maintenance and differentiation in the mammalian intestinal epithelium^1,4,5,25,26^.

Given the critical roles that AHR plays in the mammalian intestine, we hypothesized that Spineless may have evolutionarily conserved functions in the *Drosophila* midgut. We found that *spineless* is indeed upregulated in the adult midgut after bacterial infection where it limits the regenerative response and functions as a tumour suppressor in two independent models.

## Results

### Spineless limits intestinal stem cell proliferation after *Pseudomonas entomophila* infection

Given the critical functions of AHR in the mammalian intestine, we sought to determine if Spineless has similar evolutionarily conserved roles in the *Drosophila* midgut. We infected flies with the enteropathogenic bacterium *Pseudomonas entomophila* as a model of intestinal damage and regeneration. Bacterial infection induced expression of antimicrobial defence genes *DiptB, Nuox, Upd3* (Fig S1A-C). *Spineless* was expressed at low levels in the steady state intestine and was induced over 200-fold 24h following bacterial infection (Fig. 1A). Expression of the Spineless binding protein *tango* (*tgo*) remained largely unchanged after infection (Fig 1B). We analysed published RNA-sequencing data of different intestinal cell populations from Dutta and colleagues^27^ which confirmed low expression of *spineless* in the steady-state midgut and strong induction after *P. entomophila* infection (Fig. S1D). Of note, *spineless* was only induced in intestinal stem cells (ISC) and enteroblasts (EB), but not in enterocytes or enteroendocrine cells. This suggests a potential role for Spineless in the progenitor compartment following bacterial infection.

**Figure 1:**
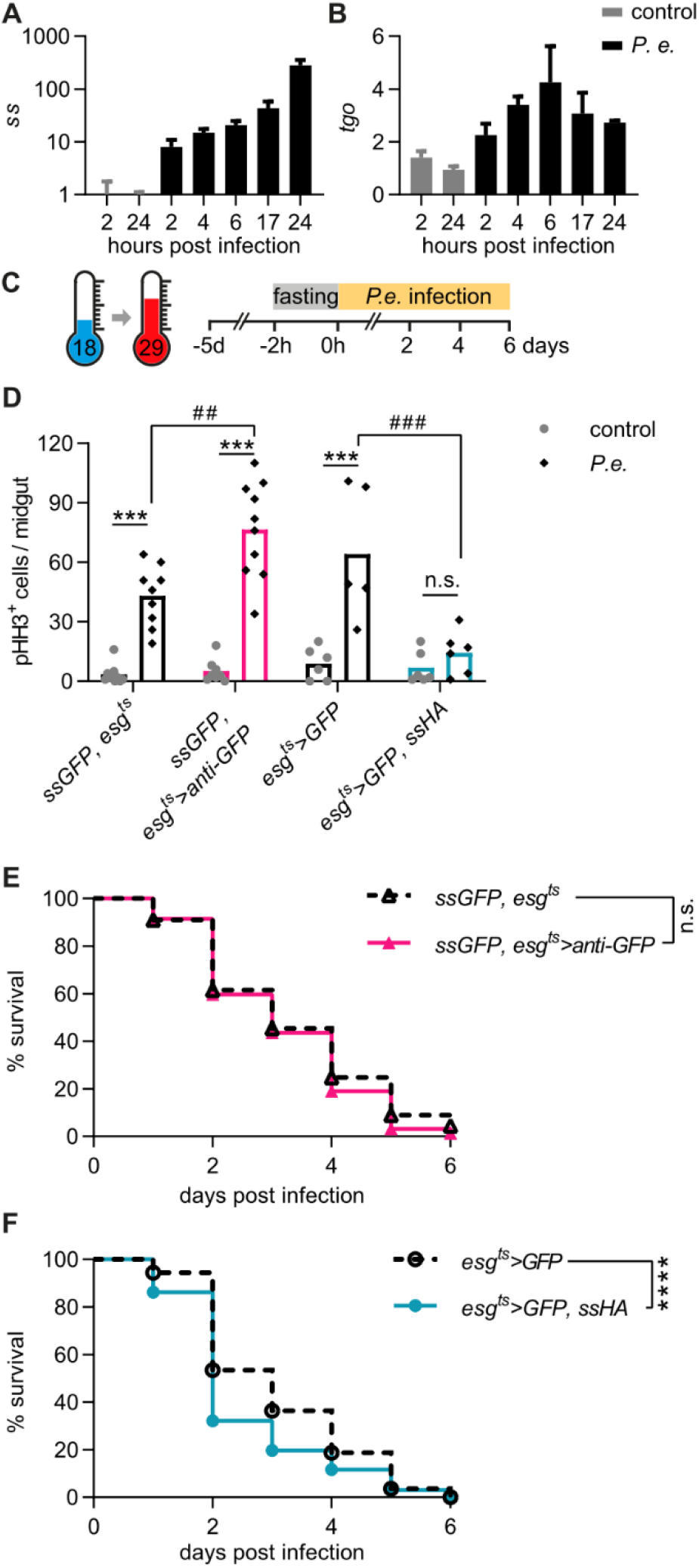
*Spineless* limits intestinal stem cell proliferation after *Pseudomonas entomophila* infection. A, B) Gene expression was determined by qPCR from isolated midguts of *w1118* flies at different timepoints after *P. entomophila* infection. Data are from one experiment with n=3 samples per timepoint. Gene expression was normalized to *Rpl32* and uninfected controls. C) Schematic of *P. entomophila* infection. D) The number of pHH3^+^ cells per midgut was quantified at 24h post *P. entomophila* infection. Data are from 3 independent experiments, n=5-10 per genotype. E, F) Survival following *P. entomophila* infection. Data are pooled from 5 independent experiments, n=230-300 flies per genotype. *P*.*e*., *Pseudomonas entomophila*.

We generated new lines to conditionally knockdown or overexpress Spineless to study its function in midgut progenitors. *ssGFP* flies were created by targeted insertion of GFP at the C-terminus of the endogenous *spineless* locus (Fig S1E). Homozygous *ssGFP* flies showed no apparent phenotype and *ssGFP* was clearly visible in the antenna imaginal disk (Fig. S1F). Flies expressing a membrane-anchored anti-GFP nanobody (*UAS-anti-GFP*) were created to prevent *ssGFP* from translocating to the nucleus (Fig. S1G). To achieve cell type-specific Spineless knockdown we utilized the GAL4/UAS system. GAL4 can be expressed under a cell type-specific promoter to induce expression of transgenes with a GAL4 binding site (UAS). We used the promoter of *rotund*, which is expressed in the imaginal disk, to induce anti-GFP. *ssGFP, rotund>anti-GFP* flies showed aristapedia and leg phenotypes typical of Spineless mutants (Fig. S1H)^7,8^. To overexpress Spineless, *UAS*-ss flies with a C-terminal 3xHA tag were generated (Fig S1I).

We made use of the *escargot* (*esg*)-*Gal4* driver and the temperature sensitive repressor *tubGal80*^*t*s^ to manipulate Spineless expression in ISC and EB of the adult midgut only. Flies were reared at 18°C and switched to 29°C 5 days prior to infection (Fig. 1C). Infection with *P. entomophila* induces epithelial damage followed by a wave of ISC proliferation that leads to epithelial regeneration^28-31^. ISC proliferation was induced in controls following *P. entomophila* infection. Spineless knockdown further increased ISC proliferation while Spineless overexpression abolished the ISC proliferative response (Fig. 1D). We hypothesized that this could reduce the ability of Spineless overexpressing flies to survive infection. While knockdown of Spineless did not affect survival following infection with *P. entomophila* (Fig. 1E), overexpression of Spineless accelerated death following infection (Fig. 1F). This suggests that overexpression of Spineless limits ISC proliferation after *P. entomophila* infection, which impairs epithelial regeneration and thereby leads to reduced survival.

### Spineless overexpression reduces survival through ISC- and EB-specific effects

In the midgut, *esg>GFP* labels ISC and EB and therefore allows quantification of these cells as well as measurement of cell size and GFP intensity. In naïve flies, Spineless overexpression increased *esg>GFP* positive cells on fluorescent images of the midgut compared to wildtype flies (Fig. 2A, Fig. S2A), suggesting that it affects intestinal progenitor dynamics. In wildtype flies, *P. entomophila* infection resulted in a noticeable increase in the number of *esg>GFP* positive cells and an increase in their cell size. Spineless overexpression blocked the increase in cell number and size after *P. entomophila* infection, in line with the lack of increased ISC proliferation measured by pHH3^+^ cell quantification (as seen in Fig. 1D).

**Figure 2:**
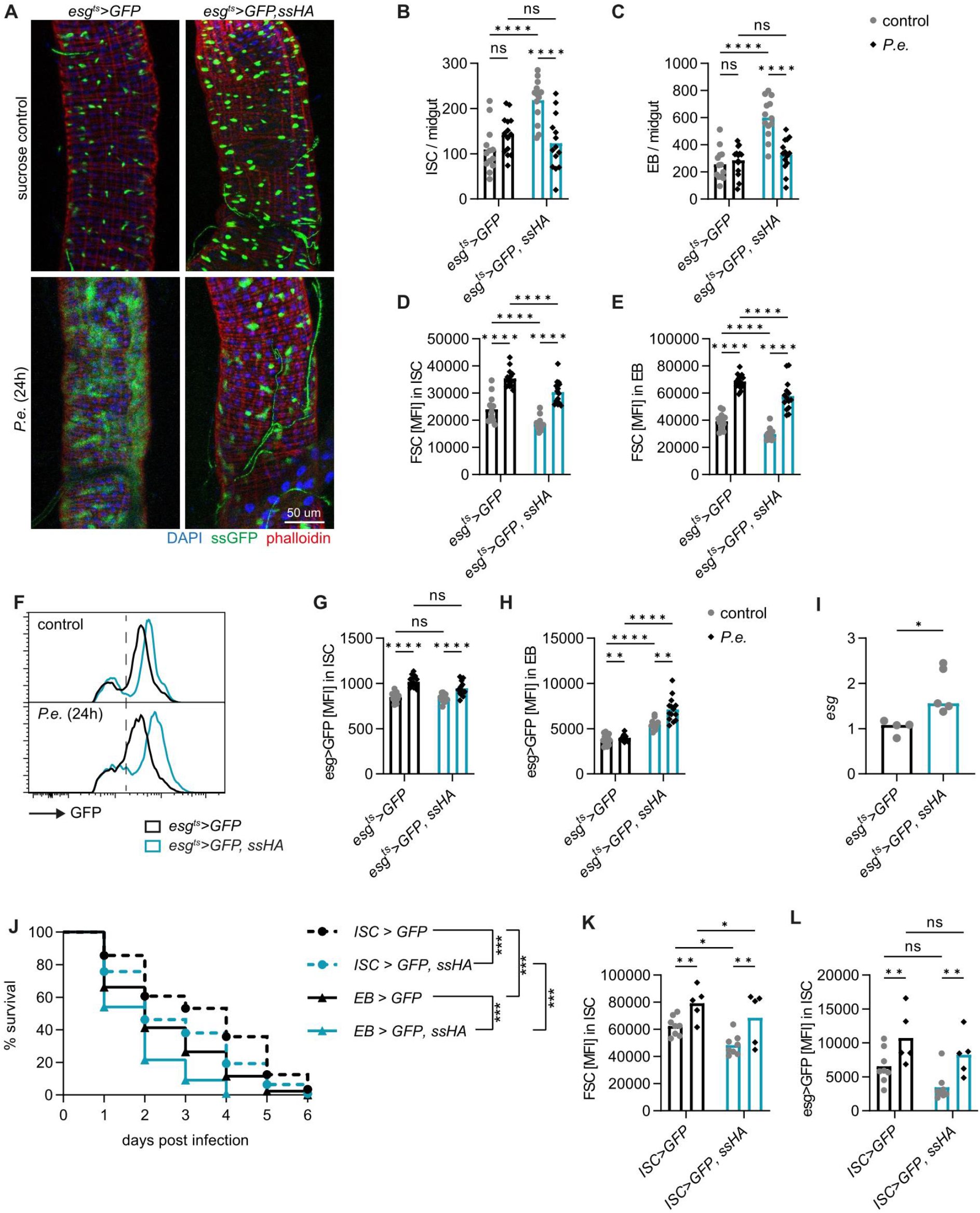
*Spineless* overexpression reduces survival through ISC- and EB-specific effects. A) Representative images of *P. entomophila* infected midguts at 24h post infection. B-H) Midguts from uninfected controls and at 24h post *P. entomophila* infection were analysed by flow cytometry. B, C) Quantification of ISC and EB numbers per midgut. D, E) Mean fluorescent intensity for FSC (cell size) in ISC and EB populations. F) Representative flow cytometry plots depicting GFP fluorescent intensity in GFP^+^ cells. G, H) Geometric mean fluorescent intensity of GFP in ISC and EB populations. B-E, G, H) Data are pooled from 2 independent experiments, n=13-14 samples per genotype. I) *Escargot* (*esg*) expression was determined by qPCR in FACS sorted EB from naïve flies and gene expression was normalized to *Rpl32*. Data are pooled from 2 independent experiments, n=4-5 per genotype. J-L) *P. entomophila* infection in flies overexpressing *spineless* specifically in ISC (*esg-Gal4, Su(H)-Gal80, tub-Gal80*^*ts*^) or in EB (*Su(H)-Gal4, tub-Gal80*^*ts*^). J) Survival following *P. entomophila* infection. Data are pooled from 2 independent experiments, n=295-339 flies per genotype. K, L) Fluorescent intensity of FSC and GFP in ISC populations from uninfected controls and at 24h post *P. entomophila* infection. Data are from one experiment with n=5-8 samples per genotype.

We then used flow cytometry to quantify the intestinal cell populations. *Esg>GFP* positive cells are readily detectable by flow cytometry after gating on live, single cells (Fig. S2B, C). EB can be distinguished from ISC based on their higher GFP fluorescent intensity and larger cell size as has been previously reported^32^ (Fig. S2D, E). We confirmed higher *Delta* expression in ISC than EB populations by qPCR from FACS sorted cells (Fig. S2F). This approach corroborated that Spineless overexpression increased ISC and EB numbers at steady state and lead to a decrease in the number of ISC and EB in infected versus naïve Spineless overexpressing flies (Fig. 2B, C). It also confirmed an increase in cell size in wildtype flies after *P. entomophila* infection and a reduction in cell size in naïve or infected flies overexpressing Spineless compared to wildtype flies (Fig. 2D, E).

Within the ISC and EB populations, *esg>GFP* fluorescence increased after infection in wildtype controls (Fig. 2F-H). Escargot was shown to maintain stemness in ISC and EB^33,34^. In EB, Escargot also enhances Notch signalling by inhibiting Amun which may promote differentiation^33^. Spineless overexpression did not affect *esg>GFP* expression in ISC (Fig. 2G), but increased fluorescence intensity in both naïve and infected EB (Fig. 2H). We confirmed that *escargot* gene expression was increased by qPCR in FACS-sorted EB from naïve flies (Fig. 2I). This suggests that Spineless directly affects *escargot* expression in EB.

We next sought to determine whether Spineless overexpression in ISC or EB was responsible for the reduced survival following *P. entomophila* infection. Overexpression of Spineless using cell type-specific drivers for either ISC or EB reduced survival after infection in both cases (Fig. 2J). Spineless overexpression in ISC reduced ISC cell size but had no significant effect on *esg>GFP* fluorescent intensity (Fig. 2K, L), in line with the results observed with the *esg*^ts^ driver. Spineless overexpression in EB did not affect EB cell size or *Su(H)>GFP* expression (Fig. S2G, H). Taken together, these results suggest that Spineless reduces survival following bacterial infection by limiting ISC proliferation and possibly by altering EB maturation through *escargot*.

### Spineless prevents intestinal tumour formation in the *Notch*^RNAi^ tumour model

Limiting the proliferation of ISC is detrimental in the context of infection-induced damage and regeneration but could be beneficial in the context of tumours. We therefore hypothesized that Spineless may prevent tumour formation in the midgut. Loss of Notch signalling (through *Notch*^RNAi^) results in the proliferation of neoplastic ISC-like cells that fail to differentiate and form multi-layered tumours^23,24,35^. Spineless knockdown had a small but significant effect on survival in the *Notch*^RNAi^ tumour model, reducing median survival from 21 to 20 days (Fig 3A, B). Tumour growth induced by *Notch*^RNAi^ can be accelerated by infecting flies with *P. entomophila*, which induces a wave of stem cell proliferation^35,36^. We therefore infected flies with a low dose of *P. entomophila* for 24h and then followed their survival. Indeed, bacterial infection reduced the median survival of tumour flies from 21 to 7 days (Fig. 3B-D). Spineless knockdown flies succumbed even faster to tumours, with a maximum survival of only 16 days compared to 37 days for control tumour flies after *P*.*entomophila* infection (Fig. 3D).

**Figure 3:**
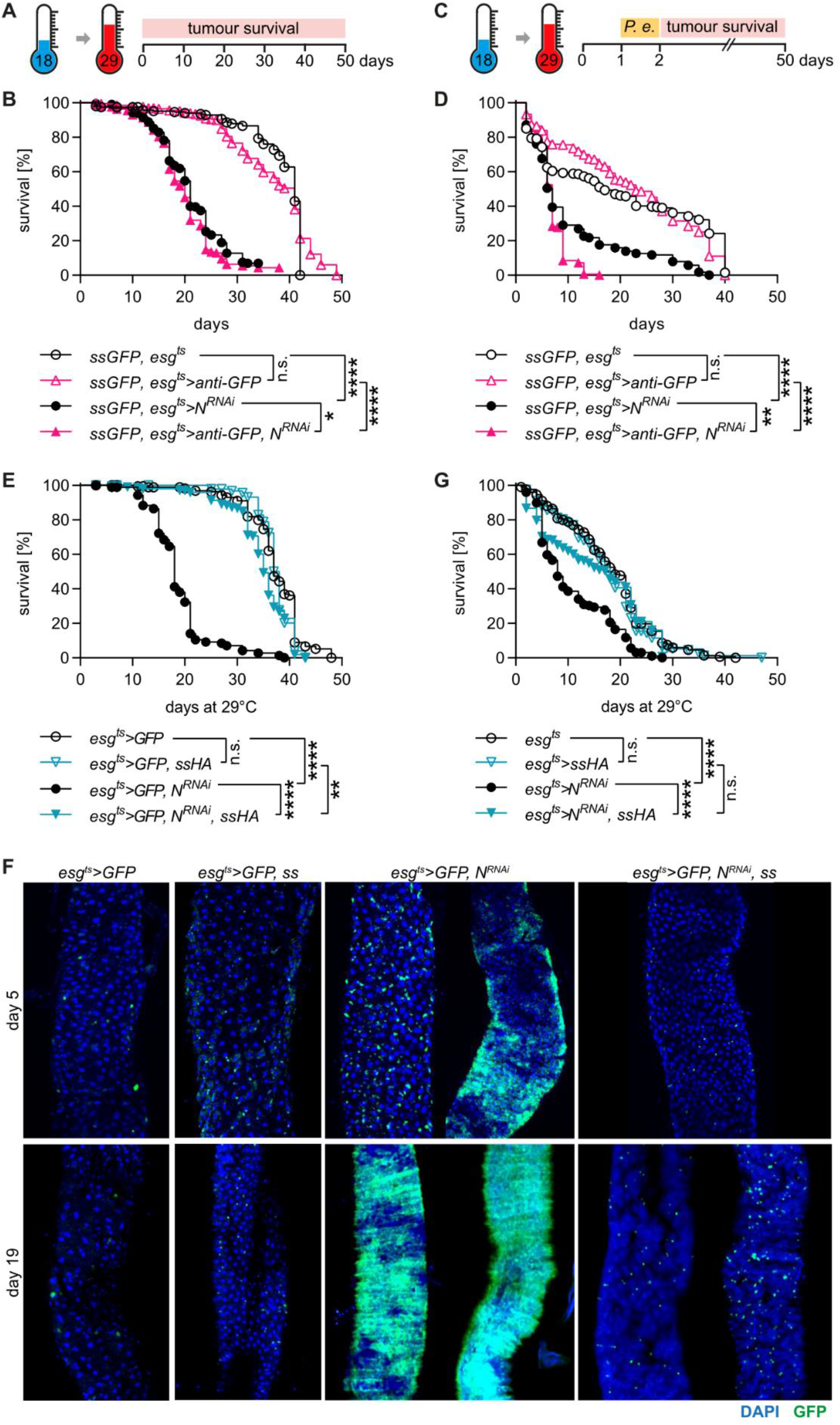
*Spineless* prevents tumour formation in the *Notch*^RNAi^ tumour model. A) Schematic of *Notch*^RNAi^ tumour model. B) Survival of *spineless* knockdown and controls in the *Notch*^RNAi^ tumour model. Data are pooled from 2 independent experiments, n=189-297 flies per genotype. C) Schematic of *Notch*^RNAi^ tumour model with 24h low-dose *P. entomophila* infection. D) Survival of *spineless* knockdown and controls in the *Notch*^RNAi^ tumour model with 24h low-dose *P. entomophila* infection. Data are from two experiments with n=166-265 flies per genotype. E) Survival of *spineless* overexpression and controls in the *Notch*^RNAi^ tumour model. Data are pooled from 2 independent experiments, n=173-202 flies per genotype. F) Representative fluorescent images of controls and *spineless* overexpressing flies at different timepoints of the *Notch*^RNAi^ tumour model. G) Survival of *spineless* overexpression and controls in the *Notch*^RNAi^ tumour model with 24h low-dose *P. entomophila* infection. Data are from two experiments with n=187-232 flies per genotype.

Spineless overexpression completely blocked the development of tumours (Fig 3E, F). The median survival was increased from 18 days to 35 days by overexpressing Spineless in the *Notch*^RNAi^ tumour model, which nearly matched the median survival of control flies (38 days). In the *Notch*^RNAi^ model increased numbers of *esg>GFP* positive cells or clonal tumours could already be seen by day 5 (Fig 3F). By day 19, the *esg>GFP* positive tumours took over most of the midgut in surviving flies. Spineless overexpressing flies showed no signs of tumour development at day 5 and at most a slight increase in the number of *esg>GFP* positive cells by day 19. Spineless overexpression also increased survival when the *Notch*^RNAi^ tumour model was combined with low-dose *P. entomophila* infection (Fig. 3G). Thus, Spineless can block tumour development in the *Drosophila* midgut, likely by inhibiting ISC proliferation.

### Spineless delays tumour formation in the *yki*^act^ tumour model

We used a second tumour model to confirm the effect of Spineless. The transcriptional coactivator yorkie can regulate ISC proliferation during midgut epithelial regeneration^31,37^. Mutation of 3 serine phosphorylation sites to alanine leads to a constitutively active form of yorkie (*yki*^act^) that is no longer subject to control by the Hippo pathway and leads to the formation of intestinal tumours^38,39^. Following temperature shift to 29°C, *esg*^*ts*^*>yki*^act^ flies had a median survival of 7 days (Figure 4A, B). Spineless overexpression increased median survival to 31 days, while control flies without *yki*^act^ had a median survival of 37-39 days. This suggests that in this tumour model Spineless overexpression can also significantly delay tumour onset. The expansion of tumour cells in *esg*^*ts*^*>yki*^act^ flies was already visible after 2 days by microscopy but not visible in spineless overexpressing flies even by day 7 (Fig. 4C).

**Figure 4:**
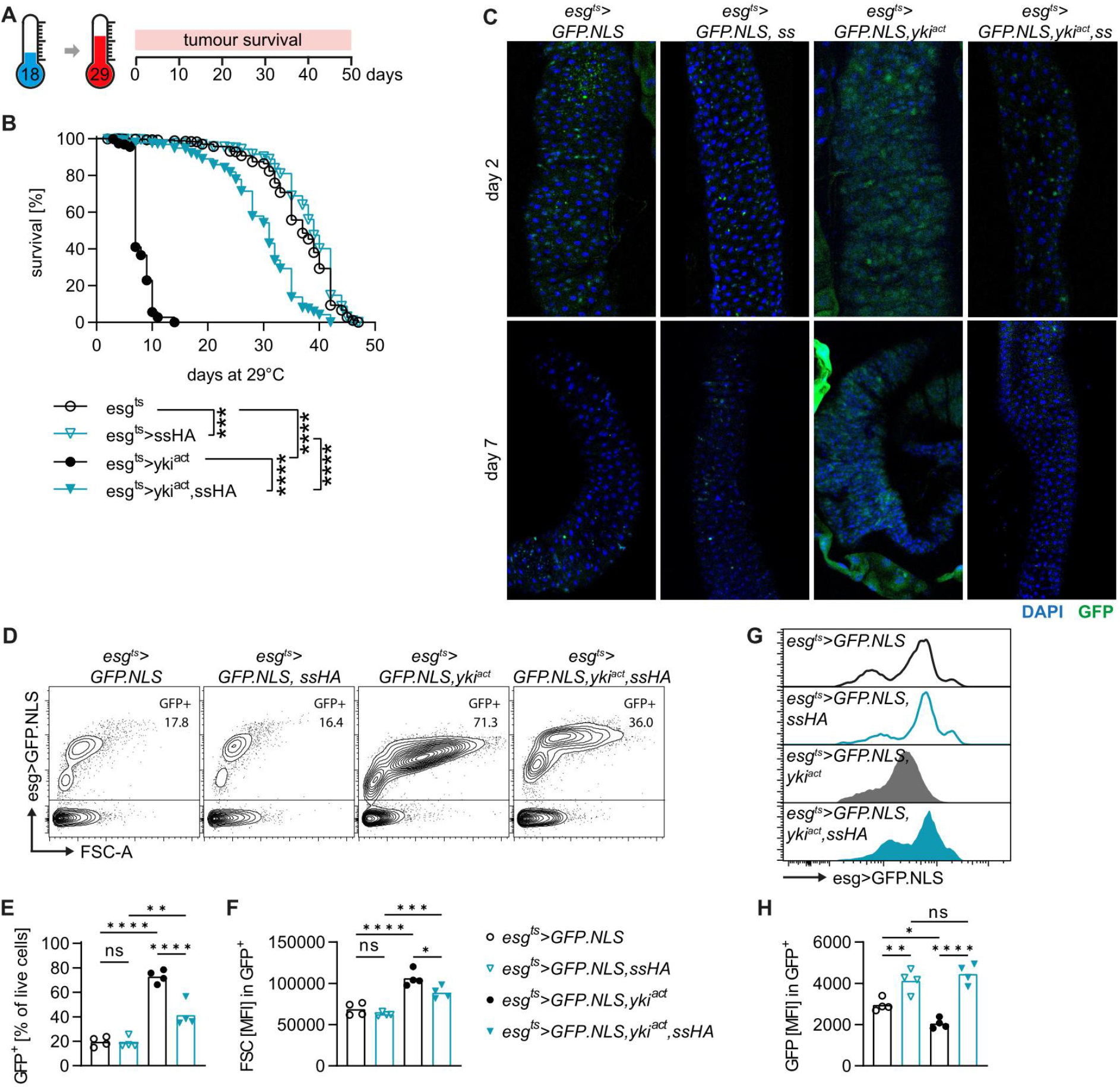
*Spineless* delays tumour formation in the *yki*^act^ tumour model. A) Schematic of *yki*^act^ tumour model. B) Survival of *spineless* overexpression and controls in the *yki*^act^ tumour model. Data are pooled from 2 independent experiments, n=409-592 flies per genotype. C) Representative fluorescent images of controls and *spineless* overexpressing flies at different timepoints after induction of the *yki*^act^ tumour model. D-H) Flow cytometric analysis of midguts from day 2 of tumour induction at 29°C. Data are from n=4 samples, each pooled of 24-30 midguts. D) Representative flow cytometry plots of GFP expressing cells as a percentage of live, single cells. E, F) Quantification of GFP^+^ cells and cell size of GFP^+^ cells. G) Representative flow cytometry plots of GFP intensity within GFP^+^ cells. I) Quantification of GFP fluorescent intensity within GFP^+^ cells.

Next, we sought to use flow cytometry to profile the GFP^+^ cells in the midgut at an early timepoint of *yki*^act^ tumour development. GFP^+^ cells were already visible 24h after temperature shift, but the distinction between ISC and EB populations was more apparent and the GFP intensity was higher after 48h (Figure S3A, B). We therefore chose the 48h timepoint. The expansion of tumour cells with EB-like fluorescent characteristics was clearly visible at this time (Fig. 4D). The frequency of GFP^+^ cells increased more than 3-fold in *esg*^*ts*^*>yki*^act^ flies and was suppressed by spineless overexpression, although not to baseline levels (Fig. 4E). Tumour cells showed an increased size (FSC) compared to GFP^+^ cells from tumour-free flies, which was partially rescued by spineless overexpression (Fig. 4F) and decreased intensity of GFP expression which was fully rescued by spineless overexpression (Fig. 4G, H). Together, these findings demonstrate that Spineless can strongly suppress or delay tumour formation in two independent models.

### Spineless affects cell metabolism, proliferation, and differentiation pathways

To understand how Spineless blocks the development of tumours, we sorted ISC and EB populations from flies with or without *yki*^act^ tumours and with or without spineless overexpression to analyse their transcriptome (Fig. S4A, B). We chose 48h post temperature shift to analyse the transition from normal to tumour cells and to be able to distinguish and isolate the ISC and EB populations by FACS. Given the differences in size and GFP intensity between the genotypes, the gates were adjusted for each genotype to best fit the populations (Fig. S4B). Principle component analysis indicated distinct clustering of ISC and EB populations (mostly along PC2) as well as genotypes (mostly along PC1) (Fig. 5A). Samples from Spineless overexpressing flies grouped away from controls in the opposite direction of *yki*^act^ tumour cells and *yki*^*act*^,*ssHA* cells clustered closer to controls than to *yki*^act^ tumour cells. This was also apparent in the number and overlap of differentially expressed genes across different comparisons in both the ISC and EB populations (Fig. S4C, D) and broadly reflects the ability of Spineless to suppress tumour growth as seen in the survival experiments. Importantly, *yki*^*act*^ and *ssHA* expression were not reduced when expressed together as compared to flies that only overexpressed *yki*^*act*^ or *ssHA* alone (Fig. 5B, C), confirming that the inhibition of tumour growth in *yki*^*act*^,*ssHA* flies is not merely the result of reduced *yki*^*act*^ expression.

**Figure 5:**
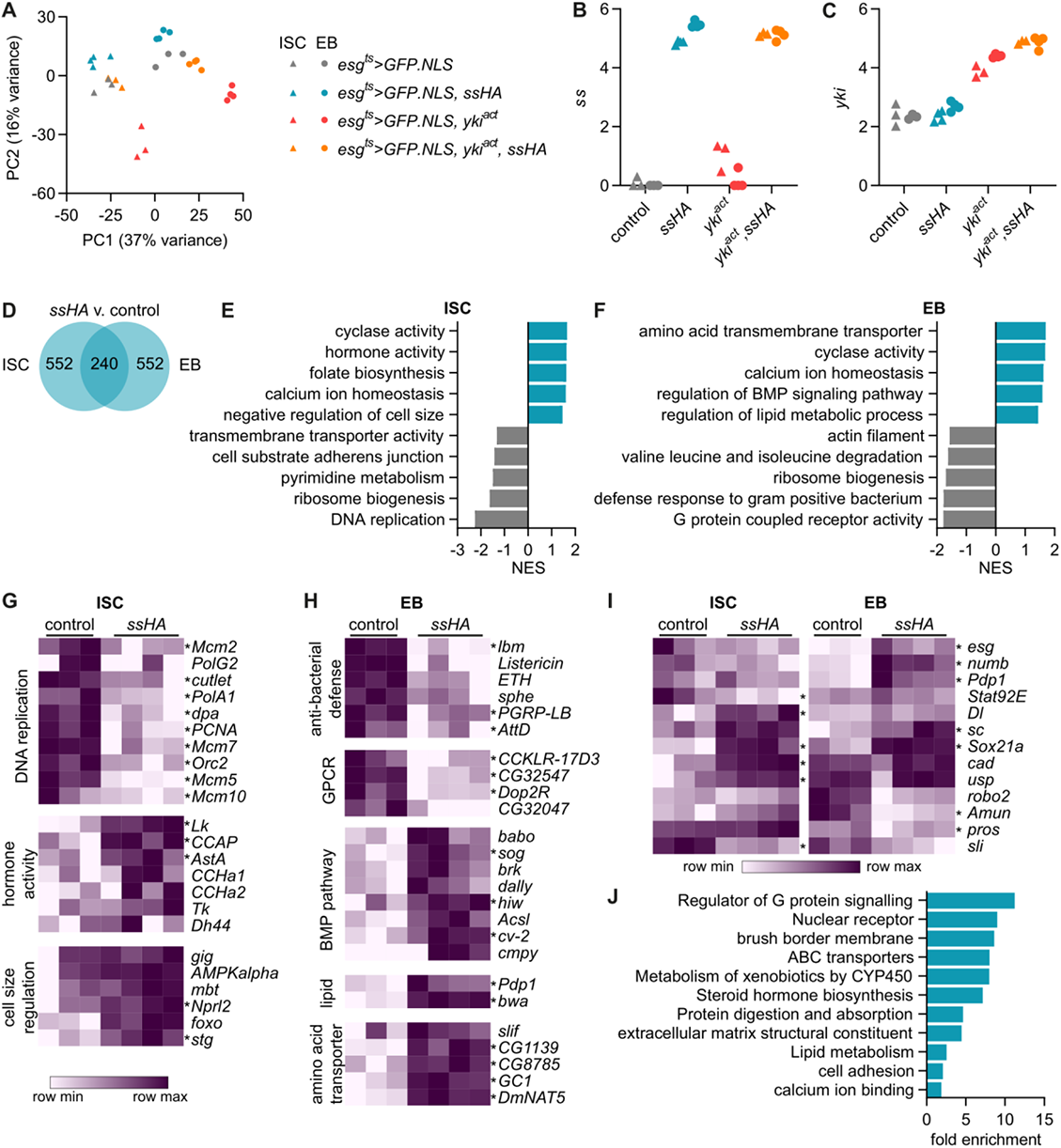
Spineless alters cell metabolism, proliferation and differentiation in midgut progenitors. A) Principle component analysis of sequenced ISC and EB samples. B, C) Expression of *ss* and *yki* in sequenced samples is depicted in log_10_(raw counts +1). D) Number of differentially expressed genes between *esg*^*ts*^*>GFP*.*NLS,ssHA* and *esg*^*ts*^*>GFP*.*NLS* samples (padj<0.05, |FC|>2). E, F) Gene set enrichment analysis comparing E) ISC and F) EB from *esg*^*ts*^*>GFP*.*NLS,ssHA* to *esg*^*ts*^*>GFP*.*NLS* samples. Selected pathways are shown, the full list is in Table S1. G), H) Examples of genes from the leading edge of enriched pathways shown in E) and F). I) Key genes involved in midgut stem cell maintenance and differentiation. G-I) Genes with significant differential expression (padj<0.05, |FC|>2) are denoted by *. J) Gene ontology analysis using DAVID of AHR-regulated mouse genes with homology to fly genes with differential expression between *esg*^*ts*^*>GFP*.*NLS,ssHA* and *esg*^*ts*^*>GFP*.*NLS* samples. Selected pathways are shown, the full list is in Table S3. Homologous genes are shown in Figure S5 and Table S2.

We first focused our attention on the gene expression changes driven by overexpression of Spineless. 792 genes were differentially expressed between Spineless overexpression and control samples in both the ISC and EB populations, with 240 genes in common (Fig. 5D, Fig. S4E). As expected, *spineless* was highly overexpressed in both cell types (Fig. S4F, G). Using gene set enrichment analysis, we found that Spineless reduced expression of genes relating to DNA replication in ISC (Fig. 5E), which correlates well with our earlier findings that Spineless suppressed cell proliferation after *P. entomophila* infection (Fig. 1). Pathways relating to DNA replication were only enriched in ISC, but not EB populations which do not proliferate (Fig. 5F). In ISC, Spineless increased expression of genes relating to hormone activity, including several hormones secreted by enteroendocrine cells (*AstA, Tk, CCha1, CCha2*)^40^ (Fig. 4G). This suggests that Spineless overexpression may result in an increase in *esg>GFP*^+^ enteroendocrine cells, a population that has previously been reported^41^. Spineless overexpression also increased expression of genes relating to negative regulation of cell size, such as *Nprl2, foxo* and *stg* (Fig. 5E, G). This is in line with our earlier findings that ISC from Spineless overexpressing flies were smaller on flow cytometry (Fig. 2D). Spineless overexpression increased expression of genes relating to calcium ion homeostasis and cyclase activity and reduced expression of genes relating to ribosome biogenesis in both ISC and EB (Fig. 4E, F).

In EB, Spineless overexpression reduced expression of anti-bacterial defence genes, GPCR activity and actin filament (Fig. 5F, H). Genes relating to lipid metabolism, amino acid transporters and the BMP pathway were increased. The BMP signalling pathway has been shown to antagonize the response to injury and return ISC to a quiescent state after injury-induced proliferation^42-44^. Genes relating to negative regulation of the BMP pathway were also enriched in Spineless overexpressing ISC, although the pathway did not reach significance (Fig. S4H, I). Inhibition of the BMP pathway by Spineless in ISC and EB may partially explain the lack of stem cell proliferation in response to infection we observed earlier.

In addition to annotated GO pathways, we also sought to directly analyse the expression of genes with known critical roles in the midgut (Fig. 5I). *Stat92E* is a key transcription factor downstream of JAK-STAT signalling. It is normally increased during midgut regeneration after injury to promote stem cell proliferation^45^. Spineless overexpressing ISC showed decreased expression of *Stat92E*, which may in part explain their reduced proliferative capacity. The mammalian genes *Cdx2* and *Rxra* are directly regulated by AHR in the intestinal epithelium^5^. Spineless increased the expression of their homologs *caudal* (*cad*) and *ultraspiracle* (*usp*) in ISC. The sequencing data confirmed increased *esg* expression in EB but not ISC from Spineless overexpressing flies, in line with our earlier flow cytometry and qPCR data (Fig. 2G-I). *Esg* has been reported to maintain stemness in ISC and EB^33,34^. In EB, Escargot also enhances Notch signalling by inhibiting Amun which may promote differentiation. Increased expression of the Notch ligand *Delta* (*Dl*) in ISC and the transcription factor *Sox21a* in ISC and EB could promote increased differentiation of EB to enterocytes^22,46-48^ and the enteroendocrine marker *pros* was decreased in EB. However, many of the differentially expressed genes promote differentiation into enteroendocrine cells. *Scute (sc)* overexpression in ISC and EB leads to an increase in Pros^+^ enteroendocrine cells^*49*^, *numb* facilitates enteroendocrine cell fate specification by limiting Notch signalling^50^, and Pdp1 is a transcription factor in enteroendocrine cells with binding sites in the promoters of hormones^51^. Spineless increased expression of *sc, numb* and *Pdp1* in EB. *Sli* encodes Slit, a ligand for the receptor Robo2. The Slit/Robo2 pathway forms part of a negative feedback loop that limits commitment to the enteroendocrine lineage^52^. We found decreased expression of s*li* in ISC and of *robo2* in EB from spineless overexpressing flies. Taken together, the RNA sequencing data suggest that spineless changes expression of a range of key factors involved in epithelial cell differentiation and may promote cell fate specification of enteroendocrine cells.

### Spineless and AHR regulate common target genes in the intestinal epithelium

To determine if AHR and Spineless had evolutionary conserved target genes in the intestinal epithelium, we generated a list of homologous genes that were regulated by AHR in mouse epithelium and by Spineless in the Drosophila midgut. This yielded 213 mouse genes and 260 homologous fly genes (Fig. S5). We then used gene ontology analysis of the mouse genes to determine which pathways were regulated by AHR/Spineless. Target genes were enriched for pathways critical for mature epthithelial cells, such as brush border maintenance, protein digestion and absorption and the extracellular matrix. Several enriched pathways were similar to those enriched in *ssHA* compared to control, such as lipid metabolism and calcium ion binding. These data suggest that AHR and Spineless control over 200 evolutionarily conserved target genes, many of which have key functions in the intestinal epithelium.

### *yki*^*act*^ induces proliferation and changes cellular metabolism in the midgut

Next, we analysed the pathways that were differentially expressed in *yki*^*act*^ tumour samples. Tumour cells clustered farthest from control cells on the PCA (Fig. 5A) and correspondingly had the highest number of differentially expressed genes, 2023 in ISC and 1859 in EB, 830 of which were in common (Fig. 6A). Tumour ISC were enriched for pathways relating to cell proliferation such as DNA replication and showed an altered metabolism with increased expression of oxidative phosphorylation and fatty acid beta oxidation pathways (Fig. 6B). Pathways relating to cell-cell adhesion, midgut development, regulation of cell growth, the extracellular matrix and antibacterial response were all downregulated in tumour ISC. In EB, tumour samples continued to upregulate pathways of cell proliferation and altered metabolism (Fig. 6C). They also upregulated genes in the SWI SNF superfamily complex such as *Iswi, Acf, HDAC1* and *Nurf-38*, which are involved in chromatin remodelling. Tumour EB downregulated pathways relating to cell-cell adhesion and the extracellular matrix similar to ISC and also reduced expression of genes involved in cilium morphogenesis, GPCR signalling and hormone activity. This shows that *yki*^*act*^ tumour cells reduced expression of genes critical for the normal function of epithelial cells in exchange for genes driving proliferation and altered metabolism.

**Figure 6:**
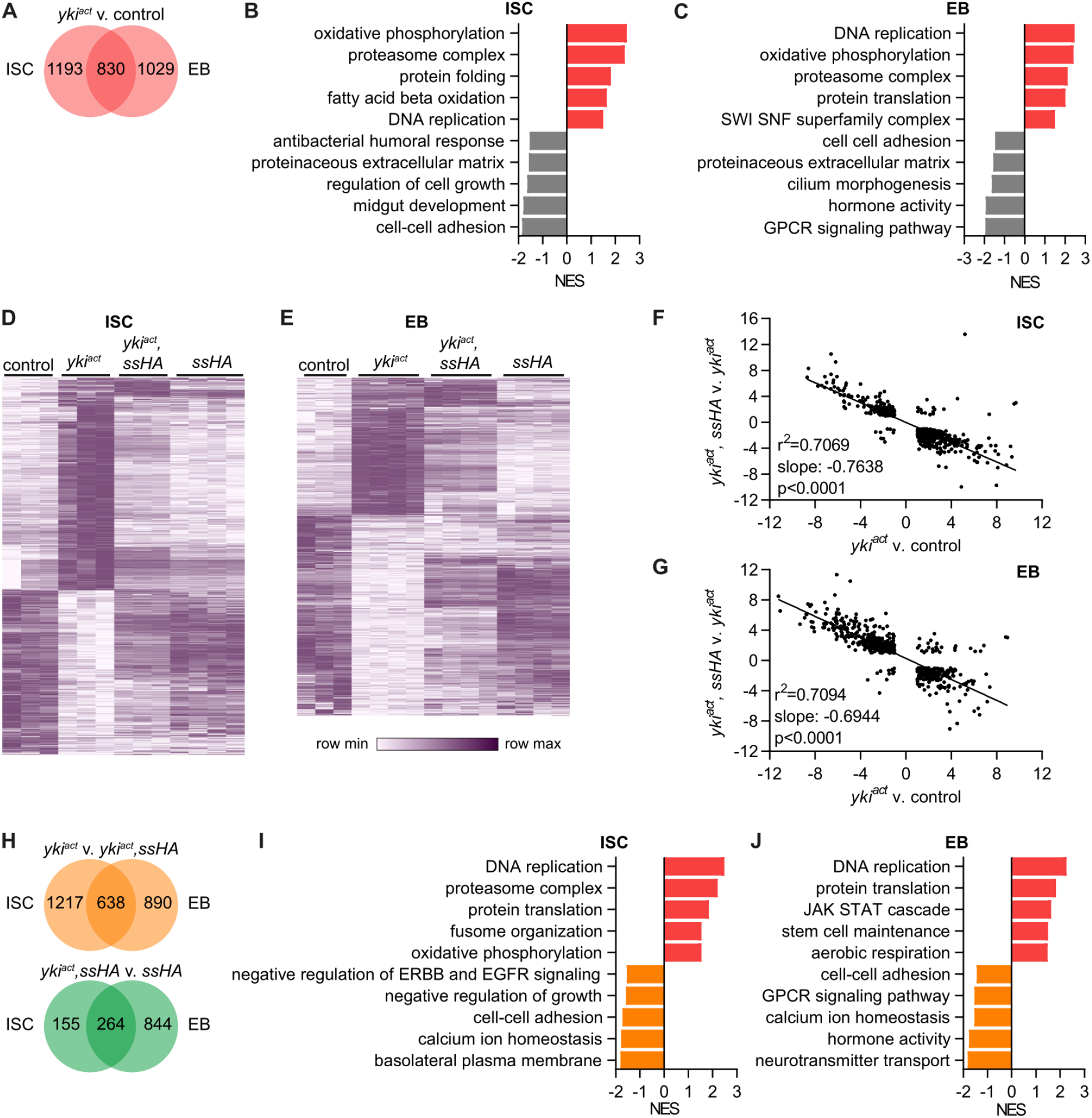
Spineless reverses effects of *yki*^*act*^ tumour on gene expression. A) Number of differentially expressed genes between *esg*^*ts*^*>GFP*.*NLS,yki*^*act*^ and *esg*^*ts*^*>GFP*.*NLS* samples (padj<0.05, |FC|>2). B, C) Gene set enrichment analysis comparing B) ISC and C) EB from *esg*^*ts*^*>GFP*.*NLS,yki*^*act*^ to *esg*^*ts*^*>GFP*.*NLS* samples. Selected pathways are shown, the full list is in Table S1. D, E) Hierarchical clustering of genes differentially expressed between *esg*^*ts*^*>GFP*.*NLS,yki*^*act*^ and *esg*^*ts*^*>GFP*.*NLS* samples. D) 2023 genes with differential expression in ISC are shown. E) 1859 genes with differential expression in EB are shown. F, G) Simple linear regression analysis of F) 946 genes in ISC and G) 741 genes in EB that are differentially expressed in both comparisons of *esg*^*ts*^*>GFP*.*NLS,yki*^*act*^ to *esg*^*ts*^*>GFP*.*NLS* and of *esg*^*ts*^*>GFP*.*NLS,yki*^*act*^,*ssHA* to *esg*^*ts*^*>GFP*.*NLS,yki*^*act*^. H) Number of differentially expressed genes between *esg*^*ts*^*>GFP*.*NLS,yki*^*act*^ to *esg*^*ts*^*>GFP*.*NLS,yki*^*act*^,*ssHA* samples or between *esg*^*ts*^*>GFP*.*NLS,yki*^*act*^,*ssHA* to *esg*^*ts*^*>GFP*.*NLS,ssHA* samples (padj<0.05, |FC|>2). I, J) Gene set enrichment analysis comparing I) ISC and J) EB from *esg*^*ts*^*>GFP*.*NLS,yki*^*act*^ to *esg*^*ts*^*>GFP*.*NLS,yki*^*act*^,*ssHA* samples. Selected pathways are shown, the full list is in Table S1.

### Spineless reverses effects of *yki*^*act*^ tumour on gene expression

From the PCA it was apparent that *yki*^*act*^ tumour samples clustered separately from controls and concurrent spineless overexpression reversed this effect so that *yki*^*act*^,*ssHA* samples clustered closer to controls. This reversal was also visible on the level of individual genes. Hierarchical clustering of all genes with differential expression in *yki*^*act*^ vs. control ISC showed that Spineless reversed the expression of most of those genes (Fig. 6D). This effect was less pronounced in EB (Fig. 6E). For genes that were differentially expressed in both comparisons (*yki*^*act*^ v. control and *yki*^*act*^,*ssHA* vs. *yki*^*act*^), the effect of *yki*^*act*^ was almost perfectly reversed by Spineless in ISC and EB (Figure 6F, G). This effect was also visible in the number of differentially expressed genes between *yki*^*act*^,*ssHA* vs. *yki*^*act*^, which were similar to the numbers between and *yki*^*act*^ tumours and controls (Fig. 6H). In contrast, there were only 419 differentially expressed genes in ISC between and *yki*^*act*^,*ssHA* vs. *ssHA*, suggesting that the tumours had a limited effect on gene expression in the presence of Spineless. In EB, the difference was much larger with 1108 genes. On the level of pathways the comparison of *yki*^*act*^ vs. *yki*^*act*^,*ssHA* was similar to that of *yki*^*act*^ vs. controls (Fig. 6I, J). Concurrent Spineless overexpression in *yki*^*act*^ tumours suppressed the cell proliferation and oxidative phosphorylation pathways and increased pathways relating to normal epithelial function such as cell-cell adhesion, regulation of growth, basolateral membrane, hormone activity and GPCR signalling. These results clearly show that Spineless can largely reverse the effects of *yki*^*act*^ on gene expression and as a result restore normal cell function and metabolism pathways in midgut progenitors.

Taken together, our results demonstrate a critical role for Spineless in limiting stem cell proliferation, promoting epithelial cell differentiation, and acting as a tumour suppressor in the adult *Drosophila* intestine (Figure 7). This shows that Spineless has functional activity in the adult intestine and adds to a growing body of evidence that Spineless fulfils roles beyond development.

**Figure 7:**
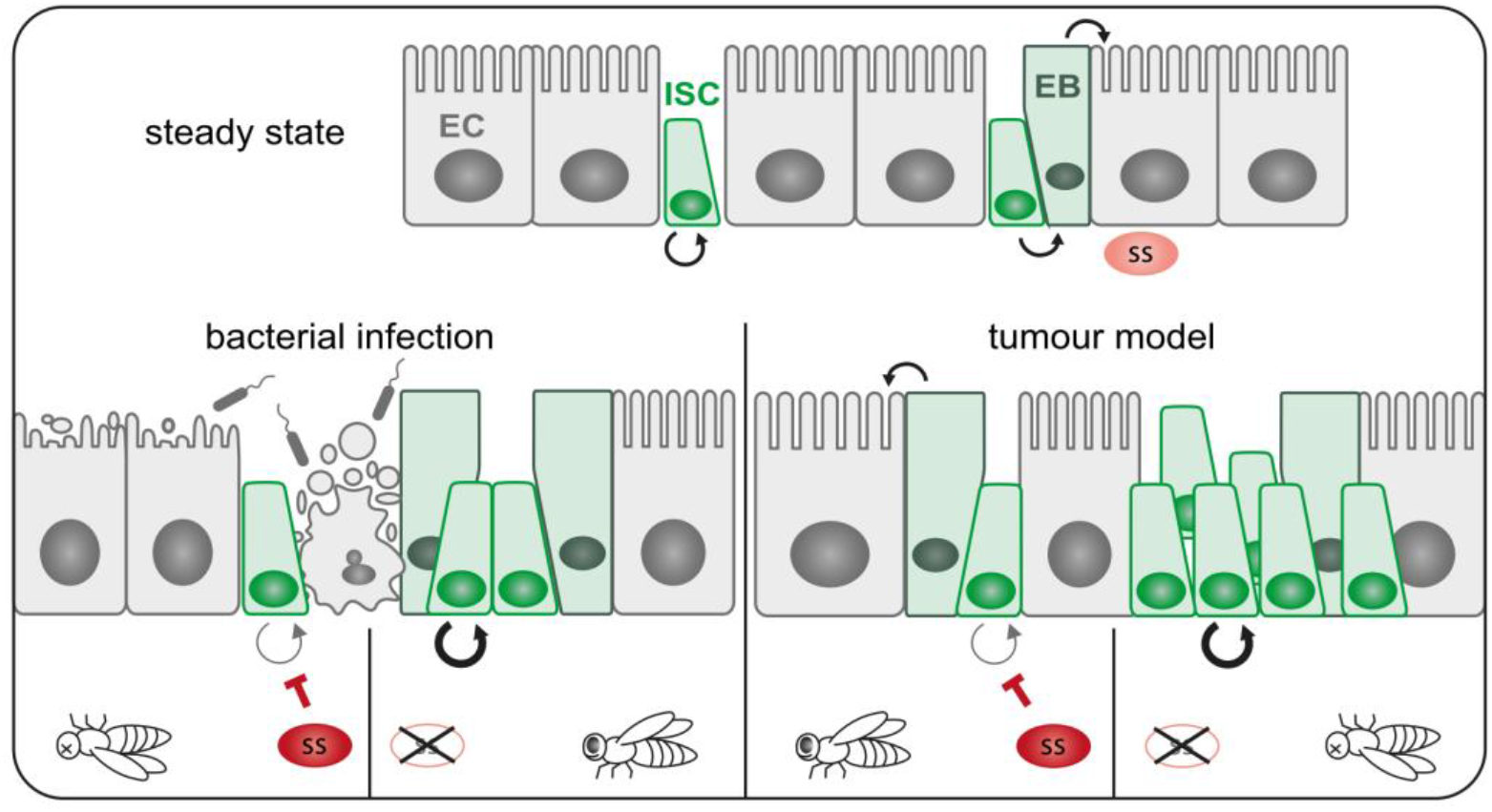
Schematic of Spineless function in the intestine. At steady state ISC undergo self-renewal in symmetric divisions or asymmetric divisions in to ISC and EB, which in turn give rise to mature enterocytes. Spineless is expressed at low levels during steady state. Bacterial infection leads to epithelial damage and stem cell proliferation to regenerate the epithelium. Spineless overexpression blocks ISC proliferation and reduces survival while spineless knockdown increases ISC proliferation. In tumour models, spineless overexpression blocks tumour growth and promotes differentiation thereby prolonging fly survival. Spineless knockdown accelerates tumour growth and decreases lifespan. EB, enteroblast; EC, enterocyte; ISC, intestinal stem cell; ss, spineless.

## Discussion

Our *Drosophila* study shows that Spineless regulates several conserved pathways in the intestine that are known to be regulated by AHR in mammals. The BMP gradient increases from crypt to villus and drives terminal differentiation of goblet cells and enterocytes in the mammalian intestine^53^. Genes in the BMP pathway were increased by Spineless overexpression in EB, suggesting that Spineless promotes differentiation of EB. Interactions between AHR and the BMP pathway have previously been suggested^54,55^, but have not been explored in the intestine. Spineless overexpression also increased expression of *cad* in ISC. Its homolog *Cdx2* is a direct target of AHR in mouse epithelial cells^5^. Both *cad* and *Cdx2* function in regionalization of the intestine^51,56^. *Cad* has also been shown to repress antimicrobial genes^57^, which could explain reduced expression of anti-bacterial defence genes following Spineless overexpression. *Cad* overexpression results in reduced ISC proliferation and epithelial regeneration after injury^58^, similar to what we observed when overexpressing Spineless during *P. entomophila* infection. Likewise, *usp*, the homolog of the direct AHR target *Rxra*^59^ was also increased by Spineless. This suggests that AHR/Spineless have evolutionarily conserved target genes with critical functions in epithelial homeostasis.

This work identified novel tumour suppressing functions for Spineless. In mice, AHR was shown to suppress colorectal cancer development in an epithelial cell-intrinsic manner^4^. We found that Spineless knockdown resulted in accelerated death from *Notch*^RNAi^ tumours and Spineless overexpression delayed tumour growth and drastically prolonged survival in two independent tumour models. This suggests that Spineless likely suppresses tumour growth on a fundamental level. Our results support two tumour-suppressing mechanisms for Spineless: the ability to restrain stem cell proliferation as observed following *P. entomophila* infection and to promote epithelial differentiation as highlighted by the transcriptomic data. Evidence for Spineless suppressing proliferation was also visible on the mRNA level, where it reversed *yki*^*act*^-induced pathways relating to DNA replication and cell division. Spineless largely reversed the effects of *yki*^*act*^ on gene expression and thereby restored normal cell function and metabolism in midgut progenitors. AHR was shown to directly affect transcription downstream of the Hippo pathway by restricting chromatin accessibility to Yap/Tead transcriptional targets^5^ and to interact with chromatin remodelling complexes^60^. It is possible that Spineless similarly alters chromatin accessibility to repress *yki* target genes. Previous work found that Spineless binding is associated with chromatin opening during butterfly wing metamorphosis^61^. The role of AHR in cancer in general is still debated and its effect seems to vary between tumour cells and immune cells^62,63^. In the mammalian intestine, several studies find beneficial functions for AHR as a tumour suppressor^1^, suggesting that the *Drosophila* model system could be used to further study tumour suppressing functions of AHR/Spineless.

Others have shown that Spineless may affect movement, the oxidative stress response, or long-term memory formation in adult flies^64,65^. Our work adds to this growing body of evidence that Spineless fulfils roles beyond development in adult *Drosophila*. In a study by Sonowal and colleagues *Drosophila* healthspan was increased by indoles in a Spineless-dependent manner^66^. This result is surprising given that the ligand-binding domain of AHR is not conserved in Spineless and Spineless nuclear translocation is independent of dioxin^6,11^. In our study, Spineless knockdown or overexpression had no apparent effect on organismal survival in the absence of tumours.

Spineless function is most likely controlled on the transcriptional level but which factors control its expression or if there are additional protein complexes that can retain Spineless in the cytoplasm similar to AHR is not known. Ligand-dependent activation of AHR in vertebrates might offer an evolutionary advantage by allowing a rapid response and the integration of environmental signals. Despite these differences in activation, AHR and Spineless bind the same DNA sequence and we show here that they regulate evolutionarily conserved target genes and pathways critical for the maintenance of intestinal epithelial homeostasis.

## Material and Methods

### Fly Stocks and manipulation of midgut progenitor cells

The following transgenic lines were used: *rotund-Gal4, Su(H)-Gal4, Su(H)-Gal80, esg-Gal4, tub-Gal80ts, UAS*-*N*^*RNAi*^ (#GD27228, Vienna *Drosophila* Resource Centre), *UAS*-*yki*^*act*^ (w*;; *UAS*-*yki*.S111A.S168A.S250A.V5; #228817 Bloomington *Drosophila* Stock Center)^38^.

The following fly lines were generated in this study: *UAS-ssHA, ssGFP, UAS-anti-GFP*.

*Esg*^*ts*^ refers to *tub-GAL80*^*ts*^, *esg-GAL4* which was used to express transgenes in midgut ISC and EB populations. We used the following driver to limit transgene expression to ISC (*esg-Gal4, Su(H)-Gal80, tub-Gal80*^*ts*^) or EB (*Su(H)-Gal4, tub-Gal80*^*ts*^) only. Drivers were crossed to *w*^*1118*^ wildtype flies or *UAS-GFP* as control. To knockdown *spineless* in midgut progenitor cells, we used flies with *ssGFP* (homozygous), *tub-GAL80*^*ts*^, *esg-GAL4, UAS-anti-GFP*. Flies with *ssGFP* (homozygous), *tub-GAL80*^*ts*^, *esg-GAL4* without *UAS-anti-GFP* served as control.

Crosses were set up at 18°C to activate Gal80^ts^, thus restricting the expression of the Gal4-induced transgenes. Adult female offspring were selected at 0-4 days of age and shifted to 29°C to induce expression of transgenes. During incubation at 29°C, flies were transferred onto fresh food every 3-4 days.

### Genetic modification of flies

To generate flies expressing Ss fused to GFP (*ssGFP*), the *ss* locus was modified by CRISPR/Cas9-stimulated homologous recombination. DNA encoding eGFP followed by a lox-3Pax3-CHE-lox cassette was inserted just before the stop codon as described^67^. To generate *UAS-anti-GFP*, we fused the CD8 ORF (NCBI Ref.: NP_001074579) to GBP (vhhGFP4)^68^ separated by a Gly/Ser linker and inserted this new ORF in pUAST. To obtain flies allowing over-expression of HA tagged version of Ss (*UAS-ssHA*), we inserted 2 HA tags (before the stop codon) in the *ss* cDNA (isoform A)^20^. The resulting DNA was inserted in pUAST before p-element-mediated transformation.

### Bacterial infection

*Pseudomonas entomophila* (stock kindly provided by Bruno Lemaitre) was grown in LB medium at 29°C for 24h. Bacterial culture was centrifuged at 3000xg for 15 min and pellets resuspended in 5% sucrose solution for a final concentration of OD_600_=200. 1ml concentrated bacteria solution was added to filter paper placed on top of standard fly food for infection. Flies were shifted to 29°C for 5 days and starved in an empty vial for 2 hours prior to infection. Survival was recorded daily.

For tumour survival experiments with low-dose *P. entomophila* infection flies were first shifted to 29°C for 1 day, then infected with bacterial solutions concentrated to OD_600_=70 in 5% sucrose and returned to normal fly food 24h later.

### Tumour survival experiments

Crosses were set up at 18°C to activate GAL80ts, thus restricting the expression of the Gal4-induced transgenes. Adult female offspring were selected at 0-4 days of age and shifted to 29°C to induce expression of transgenes. Every 2-3 days during incubation at 29°C, flies were transferred onto fresh food and survival was recorded.

### Statistical analysis

Statistical analyses were performed using GraphPad Prism (version 9). Kaplan-Meier survival curves were plotted and analysed using log-rank (Mantel-Cox) test. A p value <0.05 was considered significant. The p values of multiple comparisons of survival curves were adjusted using the Bonferroni method. 2-way ANOVA was used to analyse grouped comparisons. All data points and ‘‘n’’ values reflect biological replicates (either from single or from pooled flies).

### Immunofluorescence staining and microscopy

*Drosophila* midguts were dissected and fixed with 4% (w/v) Formaldehyde (Thermofisher) in PBS at room temperature for 30 minutes, permeabilised with PBS 0.2% Triton x-100 (PBST) at room temperature for 30 minutes and blocked with 10% (w/v) Bovine Serum Albumin (SigmaAldrich) in PBST (PBSA) at room temperature for 30 minutes. Primary antibody rabbit anti-Phospho-histone H3 (Ser10) (Cell signalling, #9701) was diluted 1:1000 in PBSA and added at 4°C overnight. Stained tissue was washed with PBS the next day and stained with secondary antibodies donkey anti-rabbit A555 (LifeTech, A21429) 1:1000 diluted in PBSA at room temperature for 3 hours followed by 1:10000 PBS-diluted DAPI staining (5mg/mL in H_2_O, SigmaAldrich, D9542) for 10 minutes before PBS washing. Ovaries and posterior abdomen were removed, and the remaining midguts were then mounted with antifade (Thermofisher, P36934) on 21-well glass slides (1 gut/well). Images were acquired on a Zeiss LSM 710 confocal microscope and were further processed in ImageJ (FIJI, version 2.1.0). Proliferating cells were manually counted under Zeiss AxioImager M1 epifluorescence microscope using 20x objective. pHH3-positive cells were counted from 3-5 whole female midguts per experiment. Images within stacks were collected at 3-5µm z-interval, 5-7 images per stack were taken to cover the complete depth of samples acquired from R2 of the midugt^69^.

### Flow Cytometry

The cell isolation protocol was adapted from Dutta *et al*^70^. 96-well v-bottom plate was prepared with 40µL/well of digestion buffer on ice (30µL PBS, 10µL of 4mg/mL Elastase, 0.4µL of 5mg/mL DNase I). 2 midguts/well were digested at 27°C with shaking for 1h, followed by pipetting 20 times for mechanical separation. Cells were washed in wash buffer (PBS, 2mM EDTA, 0.2% BSA), centrifuged and incubated with Live/Dead nIR stain (Thermofisher) at 4°C for 30 minutes, washed in PBS and fixed with 4% PFA at room temperature for 30 minutes. Samples were washed in PBS, resuspended in wash buffer with count bright beads (Invitrogen, C36950) to determine absolute cell numbers per midgut and filtered through 40µm filters. Samples were acquired on a BD Fortessa instrument (BD Biosciences) and analysed using FlowJo v10 (TreeStar). Samples were gated on single cells using FSC-A/FSC-H and SSC-A/SSC-H and to exclude debris on FSC-A/SSC-A. Dead cells were excluded based on Live/Dead near-IR staining and autofluorescence (405nm laser, 450/50 filter) before gating on GFP^+^ cells.

### Cell sorting

Midguts were dissected and digested as described above for flow cytometry. 10-30 midguts per sample were digested in 100-300µl digestion buffer. Live cells were sorted through a 70µm nozzle on a BD Fusion instrument (BD Biosciences) using BD Diva software. ISC and EB were gated according to Fig. S2B-D. Total live GFP-negative cells (mostly enterocytes) were sorted as a control.

### RNA isolation and qPCR

Entire midguts (2 midguts per sample) or sorted cells (2000-20,000) were vortexed in TriReagent and RNA was extracted according to the manufacturers protocol, using Glycogen to precipitate the RNA pellet. RNA was reverse transcribed using the High-Capacity cDNA Reverse Transcription Kit (ThermoFisher). The cDNA served as a template for the amplification of genes of interest and housekeeping genes by real-time quantitative PCR, using TaqMan Gene Expression Assays (Applied Biosystems), universal PCR Master Mix (Applied Biosystems) and the QuantStudio 7 System (Applied Biosystems). The following primer/probes were used: *Ribosomal protein L32 (*Dm02151827_g1), *spineless* (Dm02134622_m1), *tango* (Dm02373281_s1), *DptB* (Dm01821557_g1), *unpaired 3* (Dm01844142_g1), *Dual oxidase* (Dm01800981_g1), *NADPH oxidase* (Dm01826191_g1), *Delta* (Dm02134951_m1), *escargot* (Dm01841264_s1). mRNA expression was determined using the ΔC_T_ method by normalizing to *Rpl32* gene expression. To determine expression of *spineless* isoforms, genes were amplified using Power SYBR Green PCR Master Mix (ThermoFisher). Two sets of primers were used for the ss-A isoform: forward 1 (GCGAGGAGTTGGTTCCAATG), reverse 1 (ACTGCGAGTACTGCGTGTAG), 242bp product and forward 2 (GCGAGGAGTTGGTTCCAATG), reverse 2 (CGGATGCGGATGATGGTACG), 268bp prodcut. For isoform ss-C/D the following primers were used: forward 1 (GCGAGGAGTTGGTTCCAATG), reverse 1 (CTGCTGAAGCCGATCCATTC), 395bp product and forward 2 (GCGAGGAGTTGGTTCCAATG), reverse 2 (CAAATCACCAGAGGAGCGGA), 456bp product. The primers for isoform ss-C/D also generate products of 686bp and 747bp, respectively for ss-A.

### RNA sequencing and data analysis

ISC and EB populations were sorted by FACS as described above. DAPI was used as an additional staining to exclude dead cells. Cells were gated as shown in Figure S4A, B. RNA from 5,000-30,000 cells was isolated using the Qiagen RNeasy Plus Micro Kit and eluted in 15µl water. RNA quality and concentration was analysed on a Bioanalyzer and only samples with RIN>7 were used for sequencing. NEBNext Low Input RNA libraries were prepared manually following the manufacturer’s protocol (NEBNext® Single Cell/Low Input RNA Library Prep Kit for Illumina® Instruction Manual Version 5.0_5/20, NEB #E6420L). Samples were normalized to 1ng total RNA material per library in 8μl of nuclease-free water. RNA samples underwent reverse transcription, and the resulting cDNA was amplified by 10 cycles of PCR (according to the manufacturer recommendation for 1 ng input DNA). Amplified cDNA was subjected to two consecutive bead clean-ups with a 0.6X and 0.9X ratio of SPRISelect beads [B23318; Beckman Coulter] to sample volume. cDNA was fragmented enzymatically to target an insert size of ∼200bp. Adaptors (diluted to 0.6µM) were ligated to the cDNA fragments and adaptor-ligated samples were cleaned up with SPRISelect beads (ratio: 0.8x). For the amplification of the sequencing library, 25µl of Q5 Master Mix was added, plus 10µl of a unique index (NEBNext Multiplex Oligos for Illumina [NEB #E6609]). Libraries were amplified by 8 PCR cycles. Final libraries were cleaned up with SPRISelect beads (ratio: 0.9x). The quality of the purified libraries was assessed using an Agilent D1000 ScreenTape Kit on an Agilent 4200 TapeStation. Libraries were sequenced to a depth of at least 25M reads on an Illumina NovaSeq 6000 run in 101-8-8-101 configuration.

Sequencing runs were concatenated into single gzipped fastq files. These were then aligned to genome BDGP6 using nf-core/rnaseq 3.1^71^. The resulting counts file salmon.merged.gene_counts.tsv was used to create a SummarizedExperiment (https://bioconductor.org/packages/SummarizedExperiment) which was then analysed using DESeq2^72^ to produce tables of differentially expressed genes and rnk files for Gene set enrichment analysis (GSEA). PCA plots were made using the DESeq2 function varianceStabilizingTransformation. R computations were carried out using R version 4.2.3 (2023-03-15), “Shortstop Beagle”. Datasets have been deposited to GEO under accession number GSE229388. Hierarchical clustering of genes was conducted with Morpheus (Broad Institute), using one minus pearson correlation with average linkage method. GSEA was conducted using GSEA 4.2.2 (Broad Institute) with standard settings. The gene sets were taken from gsean (https://bioconductor.org/packages/release/bioc/html/gsean.html) using the GO_dme data set and these were translated into Ensembl gene ids using the biomaRt package^73^. Leading edge analysis was used to remove overlapping pathways and to identify the underlying genes. The full list of pathways is in Table S1.

We used the Homologous Gene Database (https://ngdc.cncb.ac.cn/hgd/) to obtain a list of homologous proteins between *Drosophila* and mouse. This list was then filtered on genes regulated by AHR and Spineless. Fly genes with differential expression between *ssHA* and controls with |FC|>2 in either ISC or EB were retained. For mouse genes, we used previously published RNA sequencing date from wildtype and AHR knockout intestinal organoids (GSE133092)^5^. Mouse genes with differential expression between AHR knockout and controls with |FC|>2 in either stem cell or differentiated conditions were retained. This yielded a list of 213 mouse genes and 260 homologous fly genes that are regulated by AHR and Spineless (Table S2). Gene ontology of mouse genes was analysed using DAVID (https://david.ncifcrf.gov/) and is listed in Table S3.

## Author Contributions

M.T. and J.S. performed and analysed experiments. C.A. designed and created genetically modified flies and designed experiments. M.S. carried out computational analysis of RNA sequencing data. A.F. and Y. L. performed experiments. A.G, J.P.V. and B.S. supervised the project. N.L.D. designed, performed and analysed experiments, supervised the project and wrote the manuscript.

## Acknowledgments

This work was supported by the Francis Crick Institute, which receives its core funding from Cancer Research UK (CC2016), the UK Medical Research Council (CC2016), and the Wellcome Trust (CC2016) and a Sir Henry Wellcome Fellowship 220497/Z/20/Z to N.L. Diny as well as a Wellcome Investigator Award 210556/Z/18/Z to B. Stockinger. For the purpose of Open Access, the authors have applied a CC BY public copyright licence to any Author Accepted Manuscript version arising from this submission.

We would like to acknowledge the Science technology platforms at the Francis Crick Institute. We thank the Fly Facility, especially Joachim Kurth for help with microinjections and fly maintenance, the Flow Cytometry Facility, the Advanced Sequencing Facility and the Light Microscopy Facility. We are thankful to Robert Johnston for providing the *ss* cDNA plasmid, to Nalle Pentinmikko for discussion and reagents and to Oscar Diaz, Anke Liebert, and Murali Rao Maradana for critical reading and discussion of the manuscript.

