## Supplemental Figures for "*Drosophila* AHR limits tumor growth and stem cell proliferation in the intestine"

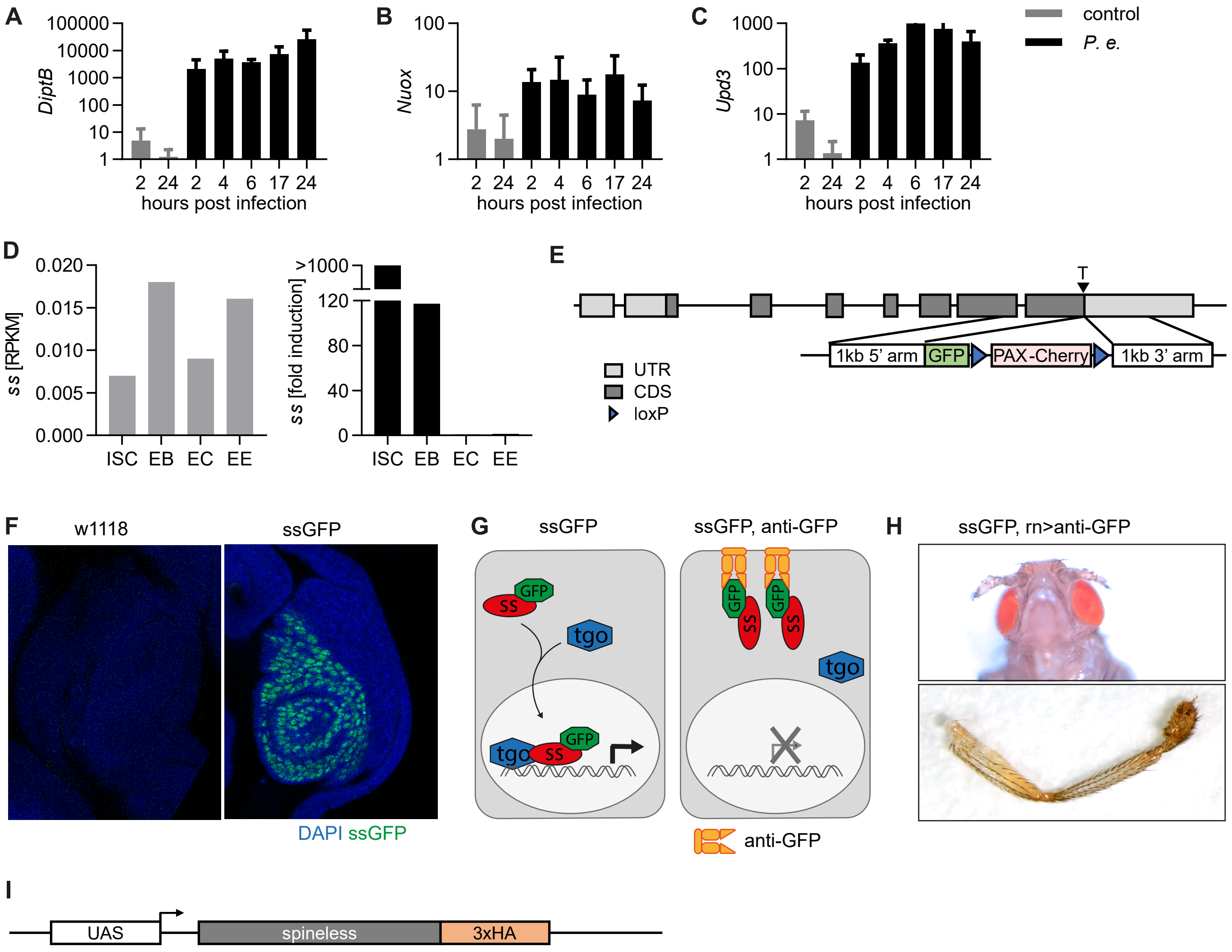


**Figure S1**
A-C) Gene expression in uninfected controls and *P. entomophila* infected w1118 flies at different timepoints after infection. Gene expression was normalized to Rpl32 and uninfected controls. Data are from one experiment with n=3 samples per timepoint.
D) RNA-Sequencing data obtained from flygutseq.buchonlab.com (Dutta *et. al.*, Cell Reports 2015). Spineless gene expression in different intesintal cell populations and induction 48h after *P. entomophila* infection is shown.
E) Schematic of the targeting contstruct to generate ssGFP flies where GFP is inserted at the C-terminus of the spineless gene.
F) ssGFP expression is visible in the antenna imaginal disk of L3 larvae from ssGFP flies. Images are from one experiment.
G) Schematic of spineless knockdown using ssGFP flies and membrane-anchored anti-GFP antibody.
H) Representative images of aristapedia and leg phenotypes typical of ss mutant flies were seen in *rn-Gal4, uas-anti-GFP, ssGFP* flies. These flies also exhibited a pharate lethal phenotype. Data are from one experiment.
I) Schematic of the transgene contstruct to generate spineless overexpressing flies.
P.e., *Pseudomonas entomophila*; RPKM, reads per kilobase per million; ISC, intestinal stem cells; EB, enteroblasts; EC, enterocytes; EE, enteroendocrine cells.

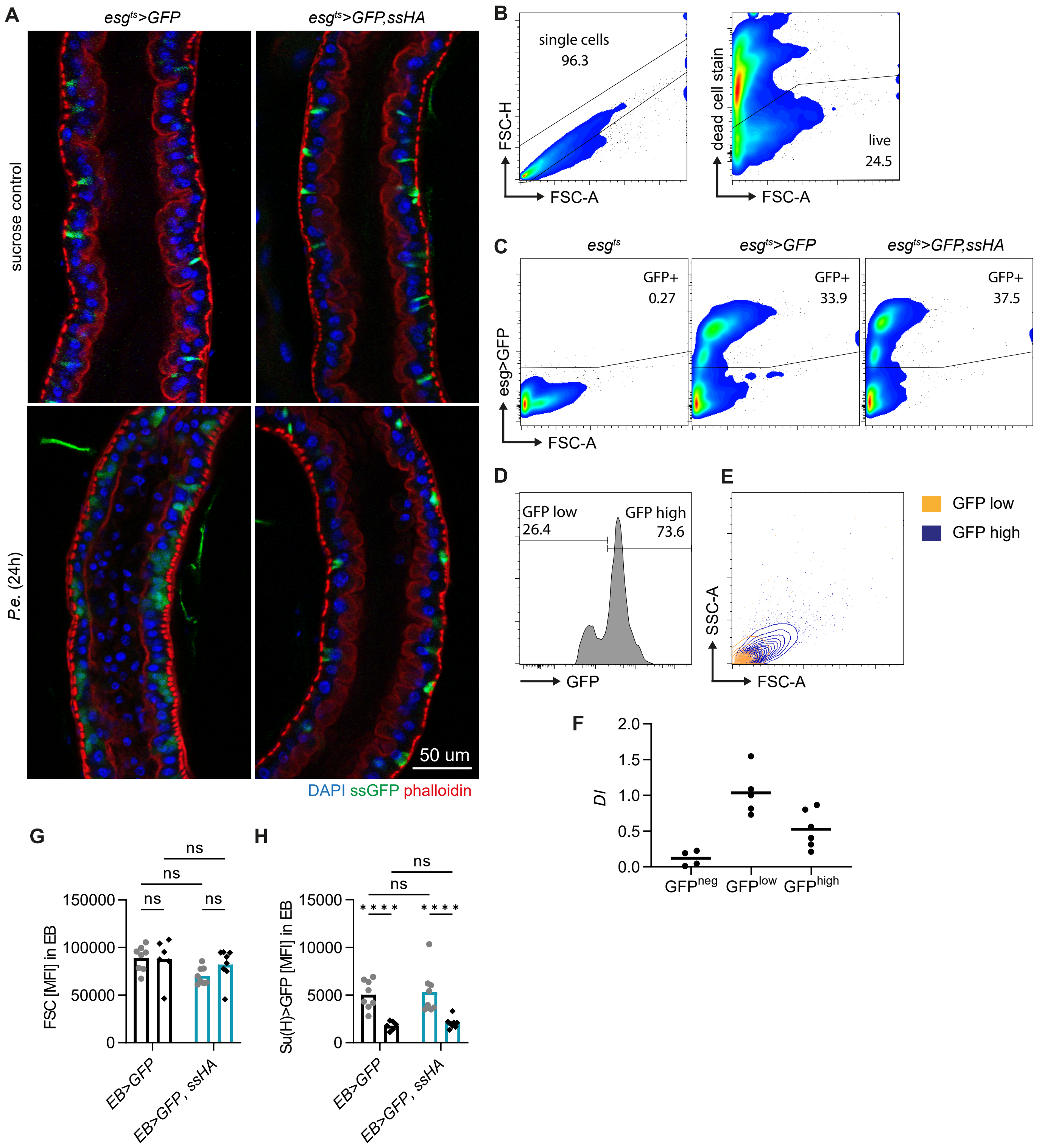


**Figure S2**

A) Fluorescent microscopy images of control and spineless overexpressing flies at 24h post *P. entomophila* infection.

B-E) Gating strategy to analyse ISC and EB populations in the midgut by flow cytometry. B) Exclusion of doublets and dead cells. C) GFP expression is shown for different genotypes. Cells were pre-gated on live, single cells as shown in B. D) $GFP^{high}$ and $GFP^{low}$ subsets amongst $GFP^{+}$ cells correspond to EB and ISC populations respectively. E) Comparison of size and granularity of the $GFP^{high}$ (EB) and $GFP^{low}$ (ISC) subsets.

F) *Delta* (*Dl*) expression in $GFP^{neg}$ (enterocytes), $GFP^{high}$ (EB), and $GFP^{low}$ (ISC) subsets was determined by qPCR in FACS-sorted cells from naive *esg$^{ts}$>GFP* flies and normalized to *Rpl32*. Data are pooled from two experiments, n=4-6 samples per cell type.

G, H) *P. entomophila* infection in flies overexpressing spineless specifically in EB (*Su(H)-Gal4, tub-Gal80ts*). Fluorescent intensity of FSC and GFP in EB populations from uninfected controls and at 24h post *P. entomophila* infection. Data are from one experiment with n=5-8 samples per genotype.

P.e., *Pseudomonas entomophila*

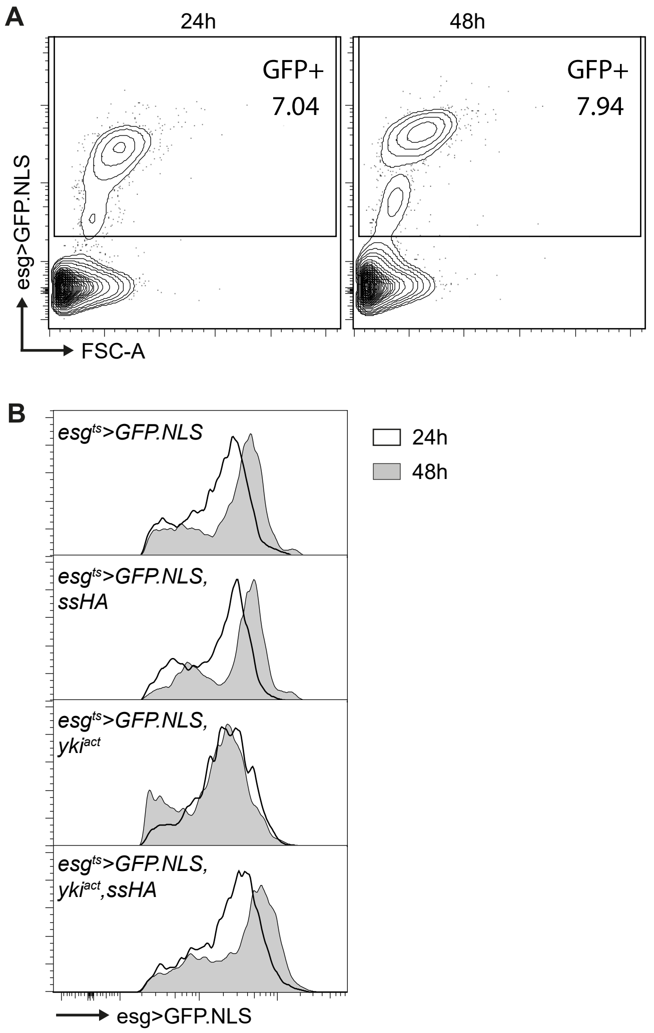


**Figure S3**

A) GFP expression in *esg*$^{ts}$>*GFP.NLS* flies at 24h and 48h after temperature shift from 18˚C to 29˚C. Cells were pre-gated on live, single cells.

B) Cells were gated on live, single, GFP$^{+}$ cells. Comparison of GFP intensity across different genotypes at 24h and 48h.

Data are from one experiment.

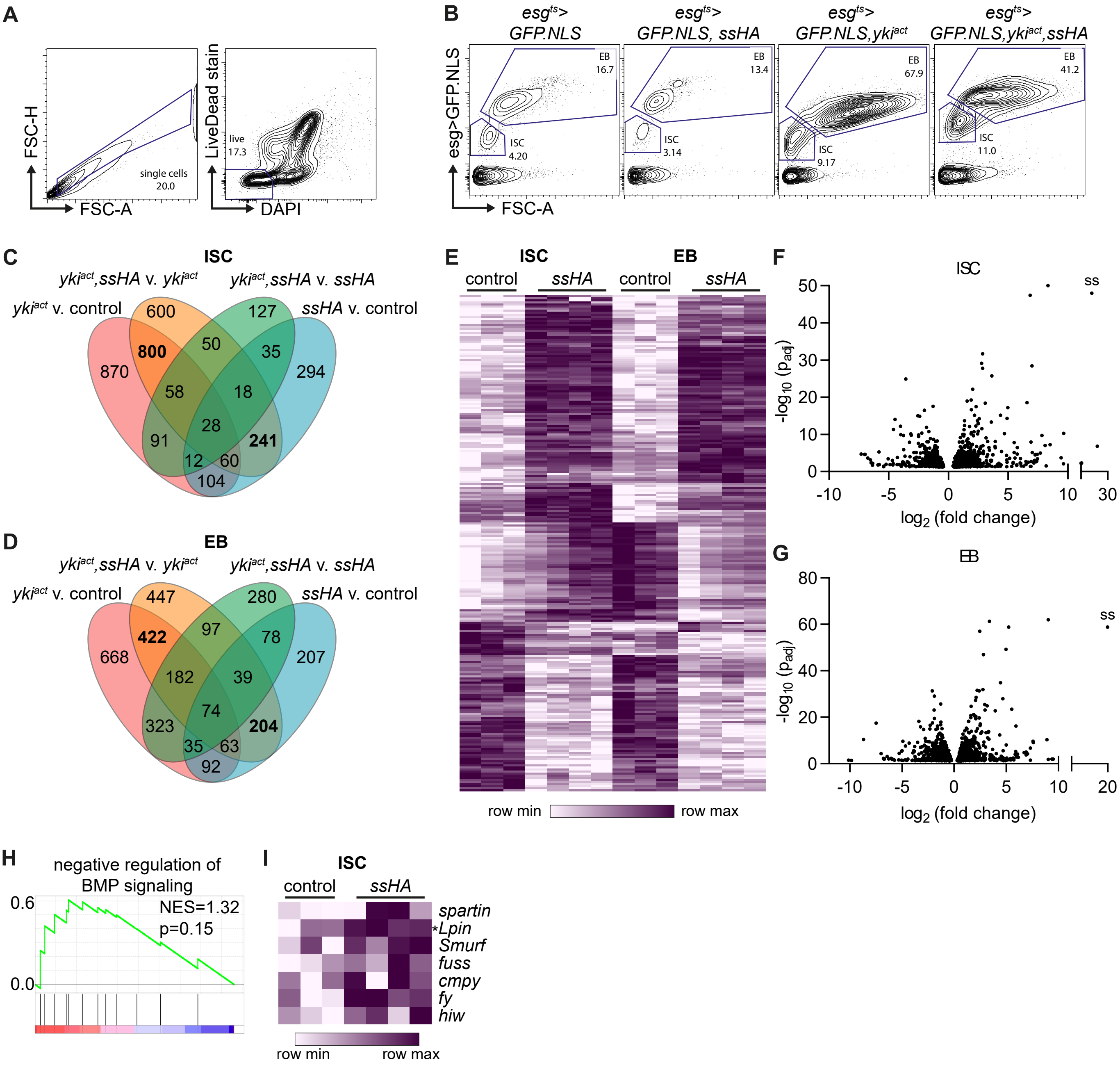


**Figure S4**

A-B) Gating of single, live cells and the ISC and EB populations in isolated midgut cells from different genotypes.

C, D) Overlap of differentially expressed genes (adjusted p value <0.05, |FC|>2). 3388 unique genes were differentially expressed across all ISC samples (C) and 3211 unique genes were differentially expressed across all EB samples (D).

E) Hierarchical clustering of 240 genes commonly differentially expressed between *esg*$^{ts}$*>GFP.NLS,ssHA* and *esg*$^{ts}$*>GFP.NLS* control samples in both ISC and EB.

F, G) Volcano plot of differentiall expressed genes between *esg*$^{ts}$*>GFP.NLS,ssHA* and *esg*$^{ts}$*>GFP.NLS* control samples in ISC (F) and EB (G).

H) Gene set enrichment analysis of the 'negative regulation of BMP signaling' pathway in ISC comparing *esg*$^{ts}$*>GFP.NLS,ssHA* to *esg*$^{ts}$*>GFP.NLS samples*.

I) Genes from the leading edge of the analysis in H).

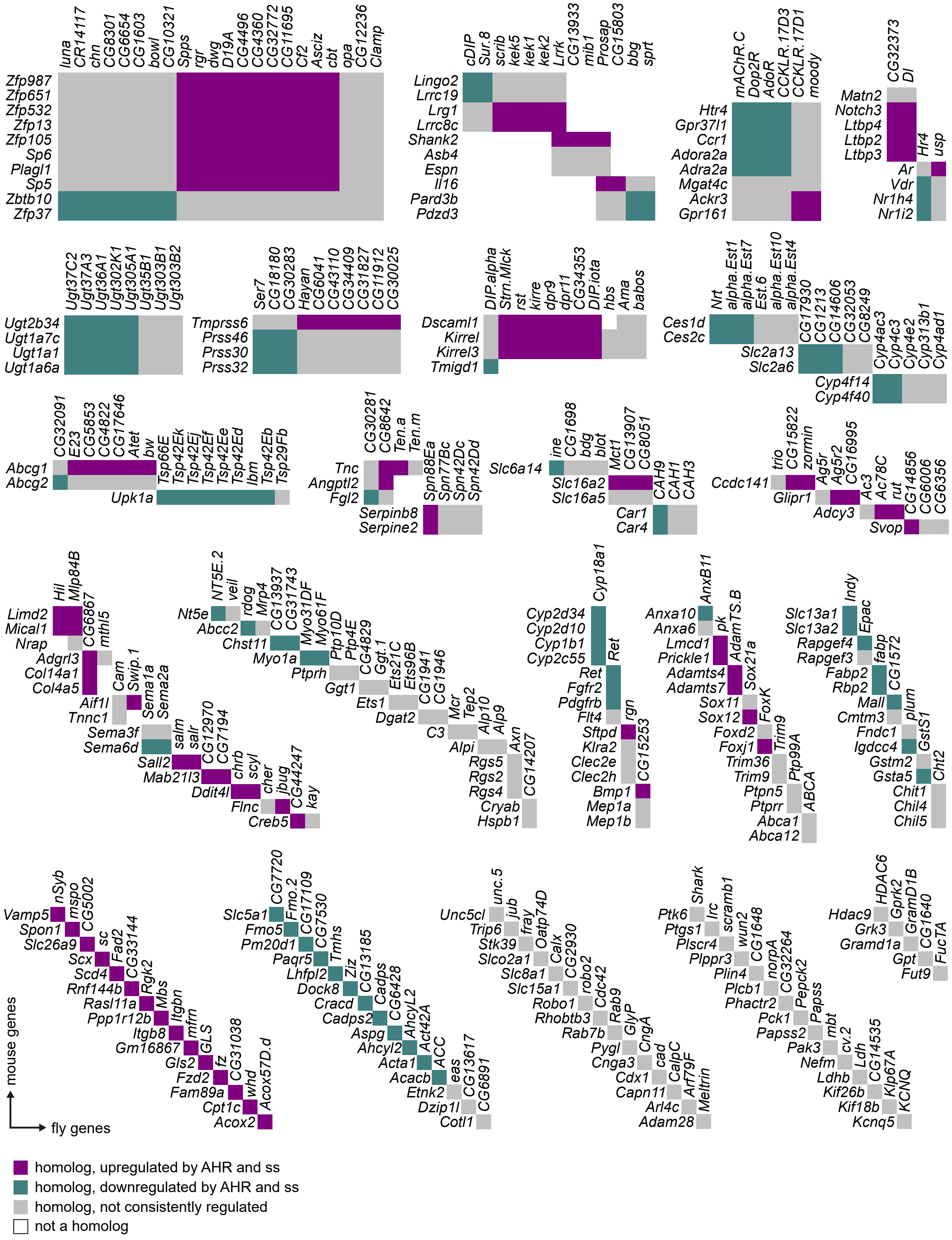


**Figure S5**

Homology between mouse genes regulated by AHR and fly genes regulated by Spineless. Beyond the homolog clusters, genes are arranged in no particular order. Data are listed in Supplemental Table 2.
